# Deoxyribozyme-based Method for Site-specific Absolute Quantification of N^6^-methyladenosine Modification Fraction

**DOI:** 10.1101/2020.01.29.925255

**Authors:** Magda Bujnowska, Jiacheng Zhang, Qing Dai, Emily M. Heideman, Jingyi Fei

## Abstract

N^6^-methyladenosine (m^6^A) is the most prevalent modified base in eukaryotic messenger RNA (mRNA) and long noncoding RNA (lncRNA). Although candidate sites for m^6^A modification are identified at the transcriptomic level, site-specific quantification methods for m^6^A modifications are still limited. Herein, we present a facile method implementing deoxyribozyme that preferentially cleaves the unmodified RNA. We leverage reverse transcription and real-time quantitative PCR along with key control experiments to quantify the absolute methylation fraction of specific m^6^A sites. We validate the accuracy of the method using synthetic RNA with controlled methylation fraction and apply our method on several endogenous sites that were previously identified in sequencing-based studies. This method provides a time and cost-effective approach for absolute quantification of the m^6^A fraction at specific loci, expanding the current toolkit for studying RNA modifications.

## INTRODUCTION

Over 100 types of RNA modifications have been identified to date. Among them, N^6^-methyladenosine (m^6^A) is most prevalent in messenger RNA (mRNA) and various long noncoding RNA (lncRNA) in higher eukaryotes (Boccaletto et al. 2018). m^6^A modifications are widely involved in post-transcriptional gene regulation. The complex and dynamic nature of m^6^A-mediated regulation enables timely responses to signaling cues and large-scale modulation of gene expression. Therefore, m^6^A has been shown to be essential for development, and associated with many human diseases (Nachtergaele and He 2018; Maity and Das 2016). The single methyl group is commonly deposited by either a methyltransferase writer complex composed of METTL3, METTL14, and WTAP (Liu et al. 2014) or by METTL16 methyltransferase (Warda et al. 2017) and is removed by either FTO (Jia et al. 2011) or ALKBH5 demethylase (Zheng et al. 2013). Through its effects on RNA secondary structure and its interactions with m^6^A binding proteins, m^6^A modifications affect essentially all known steps during an RNA’s lifetime, including alternative splicing, polyadenylation, RNA export, translation, and degradation (Kasowitz et al. 2018; Shi et al. 2019). Despite m^6^A modification having a consensus DRACH motif (D=A, G or U; R=G or A; H=A, C or U) (Meyer et al. 2012; Dominissini et al. 2012), the sub-stoichiometric nature of m^6^A modification potentially creates large compositional heterogeneity in a single RNA species, i.e., each RNA of the same species may selectively carry m^6^A modification at one or a few DRACH motifs among all (Liu et al. 2015). Being able to quantify m^6^A modification fraction at precise sites can greatly advance our current understanding of how changes in the m^6^A modification pattern (the site and fraction) are modulated by signaling cues, and are then linked to various functional consequences.

Due to its important roles, techniques have been developed and applied to detect and quantify m^6^A modification. Detection of m^6^A modification is primarily facilitated by various high-throughput sequencing-based methods utilizing antibodies and chemical crosslinking (Meyer et al. 2012; Dominissini et al. 2012; Chen et al. 2015; Linder et al. 2015). Although these sequencing-based methods can map m^6^A candidate sites at the transcriptomic level, they cannot provide the fraction of modification at each site, due to factors such as antibody binding efficiency, specificity and cross-linking reactivity (Helm and Motorin 2017). Real-time quantitative PCR (qPCR) was previously applied for locus specific detection of pseudouridine (ψ) modification though chemical labelling of ψ residue, causing a shift in the melting peak of the resulting qPCR amplicons (Lei and Yi 2017). Similar quantitative methods were recently developed for detection of m^6^A. These methods utilize enzymatic activities followed by qPCR, including differential ligation efficiency of T3 and T4 DNA ligases (Dai et al. 2007; Liu et al. 2018), differential reverse transcription activity of Tth and BstI reverse transcriptases (Harcourt et al. 2013; Wang et al. 2016), and a combination of selective elongation of DNA polymerase and ligation (Xiao et al. 2018). Although these polymer elongation and ligation-based methods are successful at modification discrimination and can report the relative m^6^A abundance change, absolute quantification using these methods were only applied on *MALAT1*, an abundant lncRNA. In addition, the potential sequence-dependence of these enzymatic activities requires caution for general applications to these methods (Potapov et al. 2018; Harada and Orgel 1993). Considering these potential pitfalls, absolute quantification using these methods would require calibration curves using fully modified and fully unmodified RNA for each target m^6^A site, which is expensive. The only available qPCR-independent method that can provide absolute quantification of m^6^A fraction site-specifically is SCARLET (site-specific cleavage and radioactive labeling followed by ligation-assisted extraction and TLC) (Liu et al. 2013). However, the sophistication of the method and its requirement for radioactive labeling prevents its broad application. Very recently, endoribonuclease digestion-based sequencing methods have been developed, which rely on selective cleavage of unmethylated A at the ACA motif (Zhang et al. 2019; Garcia-Campos et al. 2019). These approaches provide single-base resolution for identification of modifications site with relative quantitative information but are limited to m^6^A sites carrying the ACA motif, as well as regions that contain relatively sparse ACA motifs (Garcia-Campos et al. 2019). To address these challenges, we present an easy-to-implement method for quantifying m^6^A fraction at specific loci from the extracted total RNA using a deoxyribozyme (DR) with strong preference for unmethylated RNA.

## RESULTS AND DISCUSSION

### Specificity and sequence-dependence of DR cleavage efficiency

The method utilizes recently reported DR to discriminate between A and m^6^A containing RNA (Sednev et al. 2018). However, to accurately determine the modification fraction, it is important to have high cleavage efficiency for one form, and minimal for the other form to reduce the potential false positive and false negative. In addition, in order to achieve reproducible quantification, it is preferred to have discrimination efficacy less sensitive to the reaction condition. We therefore chose VMC10 DR (Fig. 1A), as it can cleave unmethylated A with reasonably high efficiency while the cleavage of m^6^A remains low even after long incubation time (Sednev et al. 2018).

**FIGURE 1.**
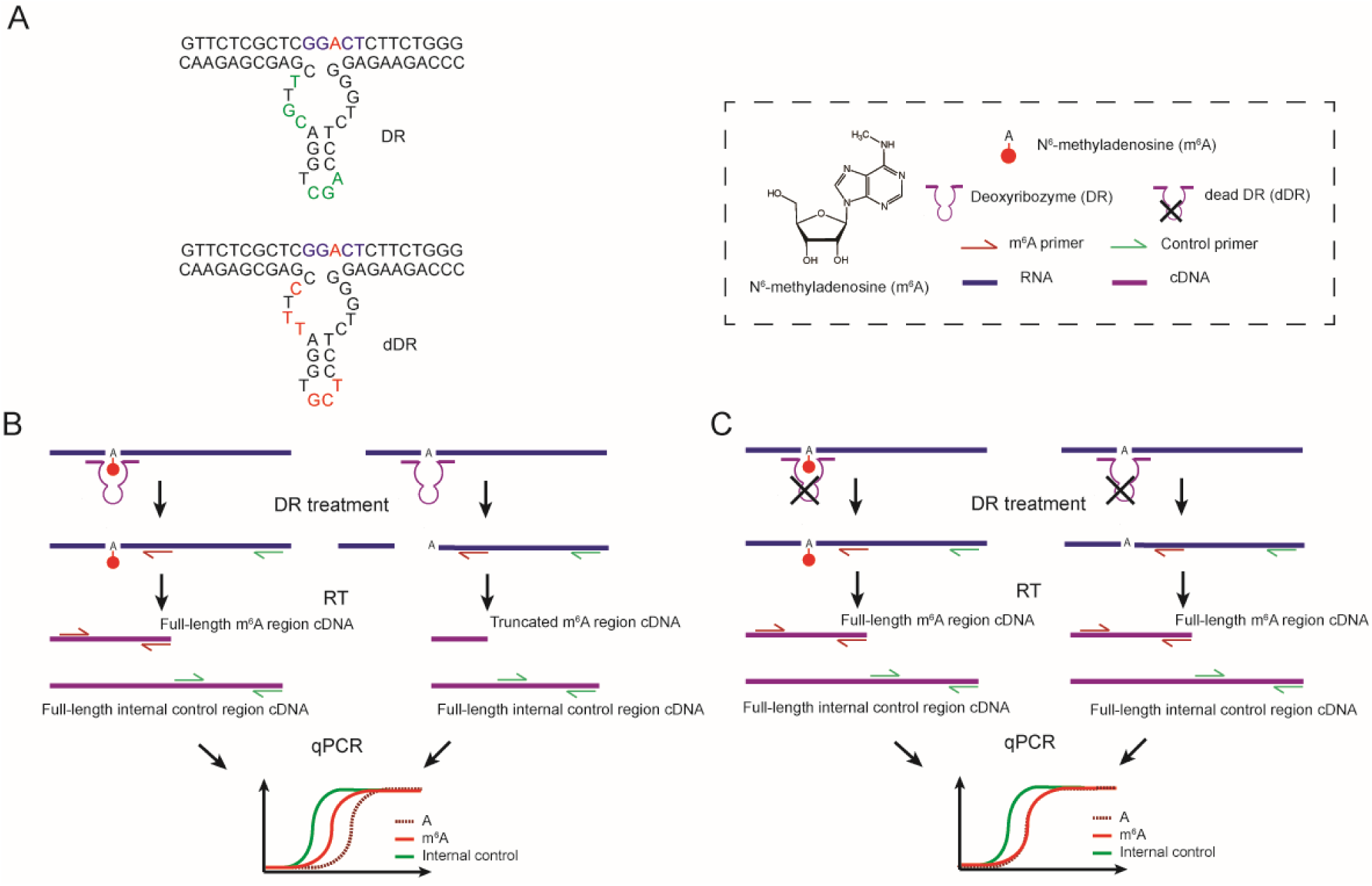
Workflow of the method. (*A*) Representative schematic of the active deoxyribozyme (DR) and the inactive deoxyribozyme (dDR). (*B*) Unmodified RNA is selectively cleaved by DR upstream of the target site, while m^6^A modified RNA remains uncleaved. The remaining uncleaved RNAs are then quantified using RT with gene-specific reverse primer and qPCR. To control for variations in RNA input, an adjacent region on target RNA is also quantified with RT and qPCR as an internal reference. (*C*) In the negative control sample, RNA is treated with a nonfunctional version of DR (dDR). Both m^6^A modified and unmodified RNA targets remain uncleaved, and are subsequently quantified with RT and qPCR.

We first verified the cleavage efficiency of DR on a variety of fully modified or unmodified sites. For this purpose, we employed a 460-nt *in vitro* transcribed RNA from a gene block sequence with only one adenine in the sequence (referred as “GB RNA” hereafter), and 35 to 41-mer synthetic RNA fragments with sequences around *MALAT1* 2515 site, *MALAT1* 2577 site, and *ACTB* 1216 site (Supplemental Table S1). Each of these targets has either m^6^A or A at the respective m^6^A sites. The RNAs were treated with corresponding 40-mer DR and subsequently analyzed by denaturing polyacrylamide gel electrophoresis (PAGE). For all the targets, the RNA fragments with unmethylated A were cleaved with high efficiency, and the cleavage efficiencies of modified RNAs were consistently below 5% (Fig. 2A-D). In addition, the cleavage efficiencies on the unmodified RNAs were sequence-dependent (Fig. 2E), ranging from 50% to 82% for our tested cases.

**FIGURE 2.**
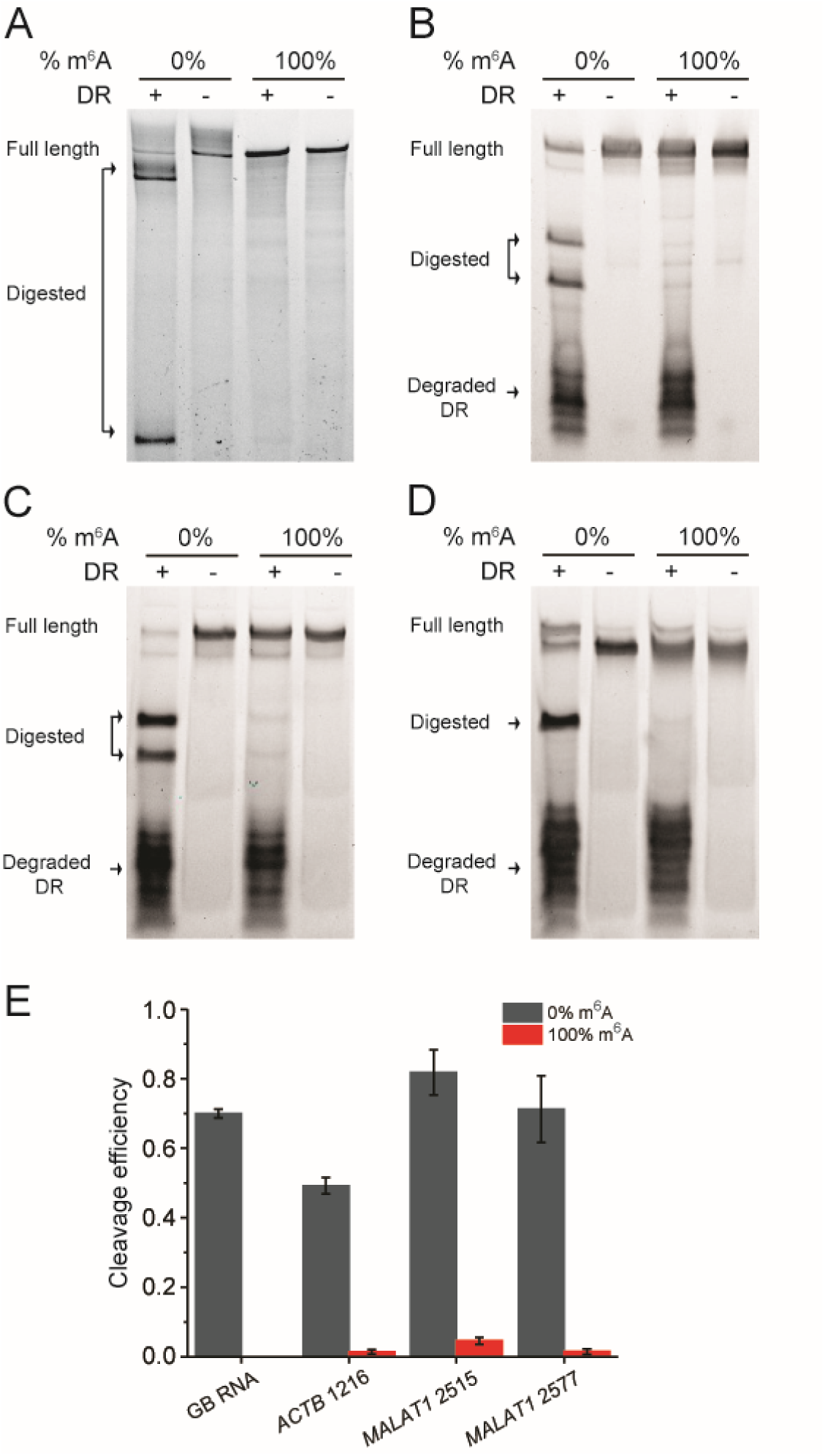
DR specifically cleaves unmodified RNAs, and its cleavage efficiency depends on sequence context around m^6^A sites. PAGE showing DR cleavage of 0% and 100% modified (*A*) GB RNA, RNA fragments containing modification site of (*B*) *ACTB* 1216, (*C*) *MALAT1* 2515 and (*D*) *MALAT1* 2577. (*E*) Bar plot of the cleavage efficiencies of m^6^A modified and unmodified target sites as quantified from PAGE. Error bars indicate mean ± s.d. for 3 to 4 independent cleavage reactions.

### Method for absolute quantification of m^6^A fraction

We designed a quantification assay using reverse transcription (RT) and qPCR. As shown in Fig. 1, DR is designed for each modification site based on a VMC10 construct. The total RNA is subjected to DR treatment during which only unmethylated RNAs upstream of the target site are cleaved. Thus, after the deoxyribozyme treatment, the amount of cleaved RNA should be inversely proportional to the methylation fraction (*Fm*) of RNA at the target site. The remaining RNA can be quantified using RT-qPCR. In order to control for the initial RNA input, we use RT-qPCR to also detect levels of adjacent uncleaved regions on the target RNA as an internal reference.

Theoretically, the modified fraction calculated from qPCR can be written as

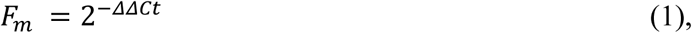

in which

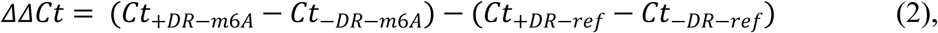

where *Ct*_+*DR-m6A*_ and *Ct*_+*DR-ref*_ are the qPCR *Ct* values at the m^6^A site and a nearby reference site in the DR treated sample, whereas *Ct*_*-DR-m6A*_ and *Ct*_*-DR-ref*_ are the *Ct* values at the m^6^A and the reference site without DR digestion.

However, the measured *ΔΔCt* only reflects the digested fraction of the RNA substrate, i.e., Eq. (1) only holds when digestion efficiency of the unmodified template is 100% and digestion efficiency of the modified template is 0%. Incomplete cleavage of unmodified A will lead to false positive, and cleavage of the m^6^A will lead to false negative. Based on the previous study and our tested cases (Fig. 2), the cleavage of VMC10 DR on m^6^A sequence is minimal, leading to insignificant error caused by false negative (Supplemental Fig. S1). In addition, it is practically difficult and expensive to generate *in vitro* purified template containing 100% modified m^6^A to account for the exactly false negative error at each m^6^A site of interest, we therefore left out the correction factor for false negative error in our final calculation. On the other hand, the false positive error can be significant due to the sequence-dependent incomplete cleavage of unmodified RNA by the DR and needs to be corrected for each m^6^A site of interest.

We therefore consider two major factors that may contribute to the false positive error due to incomplete digestion, and quantify the effect of the two factors to extract the true modification fraction: the intrinsic sequence-dependent digestion efficiency and the presence of a large amount of non-target RNAs from the total RNA extract. We define *F*_*DR*_ as a correction factor to count for the incomplete DR digestion efficiency, which has to be determined for each m^6^A target (Fig. 2). We can determine *F*_*DR*_ at each m^6^A site of interest by performing the DR digestion followed by RT-qPCR using the *in vitro* transcribed unmodified RNA:

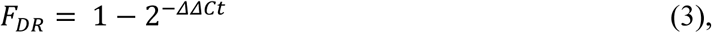

in which *ΔΔCt* is determined as in Eq. (2). We define *F*_*N*_ as the ratio of DR digestion efficiency of an RNA target in total RNA over digestion efficiency of a pure RNA target, to account for the potential drop of DR efficiency due to the presence of non-target RNAs. We can determine *F*_*N*_ by performing DR digestion using the same *in vitro* transcribed unmodified RNA mixed with total RNA, and compare with *F*_*DR*_ from Eq. (3):

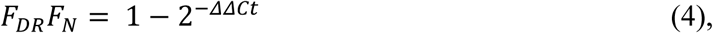

in which *ΔΔCt* is determined as in Eq. (2). With the quantification of *F*_*DR*_ and *F*_*N*_, the corrected modification fraction follows:

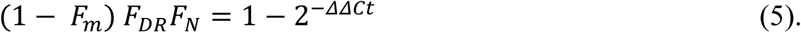

We therefore can calculate *F*_*m*_ as:

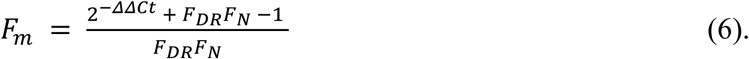

### Validation of the absolute quantification of m^6^A fraction using pure RNA

To test the feasibility of the method to quantify the m^6^A methylation fraction, we used GB RNA with methylation fractions ranging from 0% to 100%. We performed DR treatment on GB RNA and estimated the cleaved fractions by denaturing PAGE. The cleaved fraction was linearly dependent on the methylation fraction of the input RNA (Fig. 3A,B). Next, we tested whether we can use RT-qPCR to quantify the absolute methylation fraction. As this quantification is performed on *in vitro* purified RNA, only *F*_*DR*_ is needed to correct for *Fm*. Based on Eq. (3), using the 100% unmodified GB RNA, we measured *Ct*_+*DR-m6A*_ and *Ct*_+*DR-ref*_ at the m^6^A site and a nearby reference site in the DR treated sample, and *Ct*_−*DR-m6A*_ and *Ct*_*−DR-ref*_ at the m^6^A and the reference site using a negative control containing identical amount of RNA but without the DR. We found that in addition to the expected larger *Ct*_*+DR-m6A*_ compared to *Ct*_−*DR-m6A*_, there was a consistent difference between *Ct*_+*DR-ref*_ and *Ct*_−*DR-ref*_. We speculated that this difference in *Ct* values at the reference site might be due to changes in the RNA secondary structure upon DR binding that can affect RT efficiency. To create a more accurate negative control, we designed a non-functional version of DR (“dead” DR or dDR) (Fig. 1A,C), which has mutations in the AGC triplet, CG dinucleotide, and position 19 important for the catalytic activity of 8-17 family of enzymes (Santoro and Joyce 1997). We tested the activity of dDR on multiple targets, for all of which digestion of RNA was undetectable (Supplemental Fig. S2; Supplemental Fig. S3). Indeed, using the dDR treated RNA as a negative control, difference between *Ct*_+*DR-ref*_ and *Ct*_−*DR-ref*_ was eliminated (Supplemental Fig. S4). Based on Eq. (3), we determined *F*_*DR*_ of the synthetic RNA to be 0.49 ± 0.08 (mean ± s.d.). With the *F*_*DR*_ correction, we showed that the estimated *Fm* correlated well with the input m^6^A methylation fractions (Fig. 3C; Supplemental Fig. S5).

**FIGURE 3.**
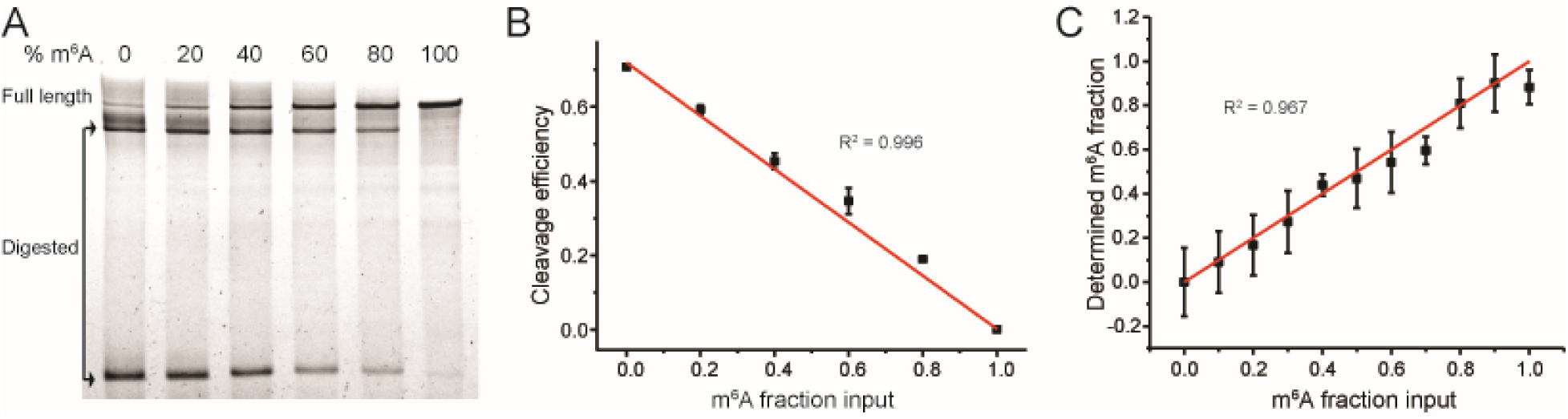
Validation of the method for absolute quantification of m^6^A fraction. GB RNA containing varied m^6^A fractions is used as a model system. (*A*) PAGE showing DR cleavage fraction of the GB RNA. (*B*) Linear relationship between the input m^6^A fraction and the cleavage fraction of RNA by DR as quantified from the PAGE gel in (*A*). Error bars indicate mean ± s.d. for 3 biological replicates. (*C*) Estimated modification fraction as a function of input m^6^A fraction for the GB RNA. Error bars indicate mean ± s.d. for at least 3 biological replicates.

### DR cleavage efficiency in presence of nonspecific RNAs

Next, we evaluated how the presence of total RNA affects the cleavage efficiency of the DR. The presence of the large amount of non-specific RNAs may compete for DR binding, consequently decreasing its cleavage efficiency at the target site in total RNA as opposed to purified RNA. We accounted for this potential decrease in efficiency with *F*_*N*_ correction factor, which we measured using three RNA transcripts: (1) the GB RNA used above, which is naturally missing in total RNA; (2) a *PLAC2* RNA fragment containing two target sites, which is of low abundance in HeLa cell line; and (3) an unmethylated A site in the endogenous *ACTB* mRNA. A1165 site on *ACTB* mRNA was chosen as the unmethylated A site because it was not detected in the sequencing-based studies (Liu et al. 2014; Ke et al. 2017), nor contains the DRACH consensus motif. *F*_*DR*_ of these three RNAs were measured with *in vitro* transcribed RNAs based on Eq. (3) (Fig. 4A,B). Then the *in vitro* transcribed GB RNA and the *PLAC2* RNA fragment were spiked into the total RNA respectively to determine *F*_*N*_ based on Eq. (4). For the unmethylated A site in the *ACTB* mRNA, *F*_*N*_ was determined by measuring the total RNA directly. In order to increase the binding specificity of DR, we also compared a 60-mer DR and a 40-mer DR. We found that *F*_*DR*_ values of 60-mer DR were higher than those of 40-mer DR (Supplemental Fig. S2), likely due to a higher hybridization efficiency by 60-mer DR. *F*_*N*_ values were consistently high for all tested RNAs, with the lowest *F*_*N*_ values being 0.78 ± 0.02 for 40-mer DR and 0.93 ± 0.02 for 60-mer DR, demonstrating that ability of our method to quantify m^6^A status should not be compromised by the presence of total RNA, and that 60-mer can slightly outperform 40-mer DR (Fig. 4C). Overall, the average *F*_*N*_ values were determined to be 0.94 ± 0.1 for 40-mer DR and 0.98 ± 0.05 for 60-mer DR. Therefore, we can simplify Eq. (6) to be

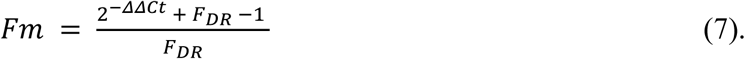

**FIGURE 4.**
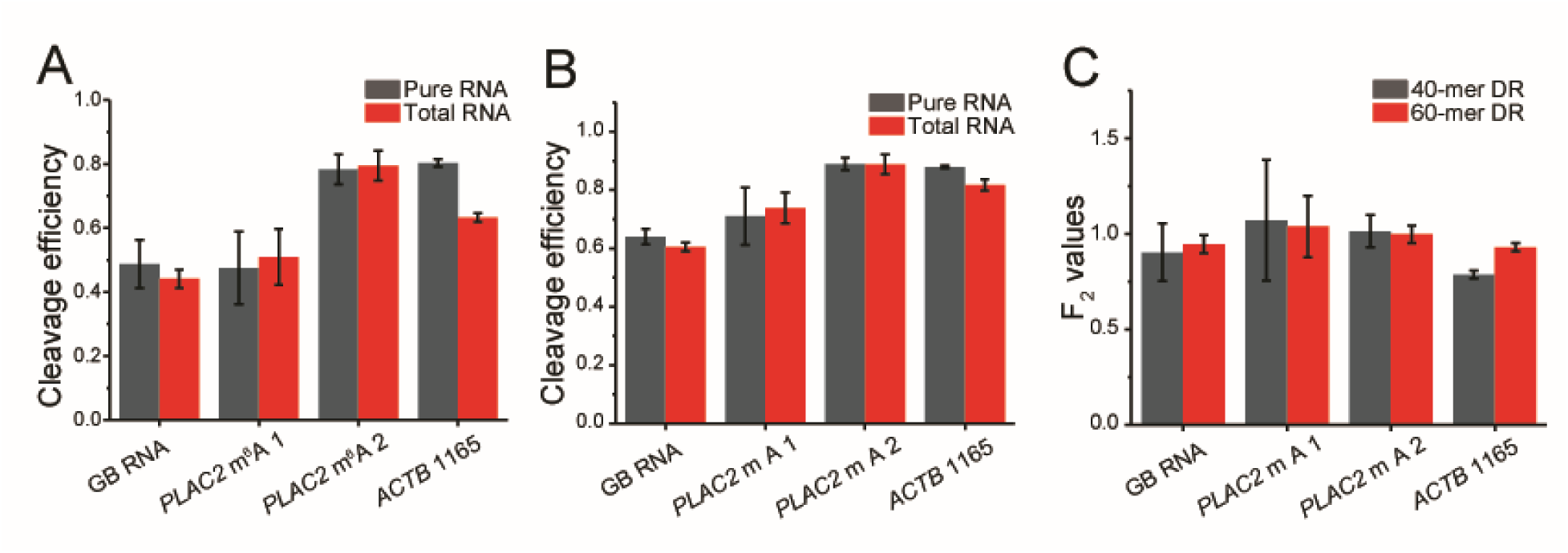
The cleavage efficiency of DR is not compromised by the presence of total RNA. The cleavage efficiencies of the GB RNA, two m^6^A sites in *PLAC2*, and *ACTB* 1165 by (*A*) 40-mer DR and (*B*) 60-mer DR in presence and absence of total RNA are determined by RT and qPCR. (*C*) F_N_ correction values for the GB RNA, two m^6^A sites in *PLAC2*, and *ACTB* 1165 for 40-mer and 60-mer DRs as determined from cleavage efficiencies in (*A*) and (*B*). All error bars report mean ± s.d. for 3 biological replicates.

### Quantification of m^6^A fraction of endogenous sites

Having developed and validated our method, we applied it to determine the methylation fraction of several endogenous sites that were identified as potential m^6^A sites by RNA sequencing from more than one study: *MALAT1* 2515 (chr11 65500276), 2577 (chr11 65500338), and 2611 (chr11 65500372), *ACTB* 1216 (chr7 5527743), *LY6K* 1171 (chr8 142703380), *MCM5* 2367 (chr22 35424323), *SEC11A* 1120 (chr15 84669674), *INCENP* 912 (chr11 62130275), 967 (chr11 62130330), and 1060 (chr11 62130423), *LMO7* 2822 (chr13 75821377), and *MRPL20* 549 (chr1 1402080) (The genome position based on GRCh38.p13 Primary Assembly of m^6^A site is indicated in parenthesis) (Liu et al. 2014; Ke et al. 2017). The selected RNAs vary from low to high abundance in Hela cells, and some of them contain more than one modification sites. To apply our method, a DR and a dDR were designed for each site. Due to the higher *F*_*DR*_ and *F*_*N*_ values with 60-mer DR, we chose to use the 60-mer DR for all endogenous RNAs. For each target site, we first generated *in vitro* transcribed RNAs containing the m^6^A sites of interest, and performed DR digestion on these *in vitro* transcribed unmethylated RNAs to get *F*_*DR*_ for each site. The *F*_*DR*_ values were all greater than 0.49 and again varied among different RNAs (Fig. 5A; Supplemental Fig. S3).

**FIGURE 5.**
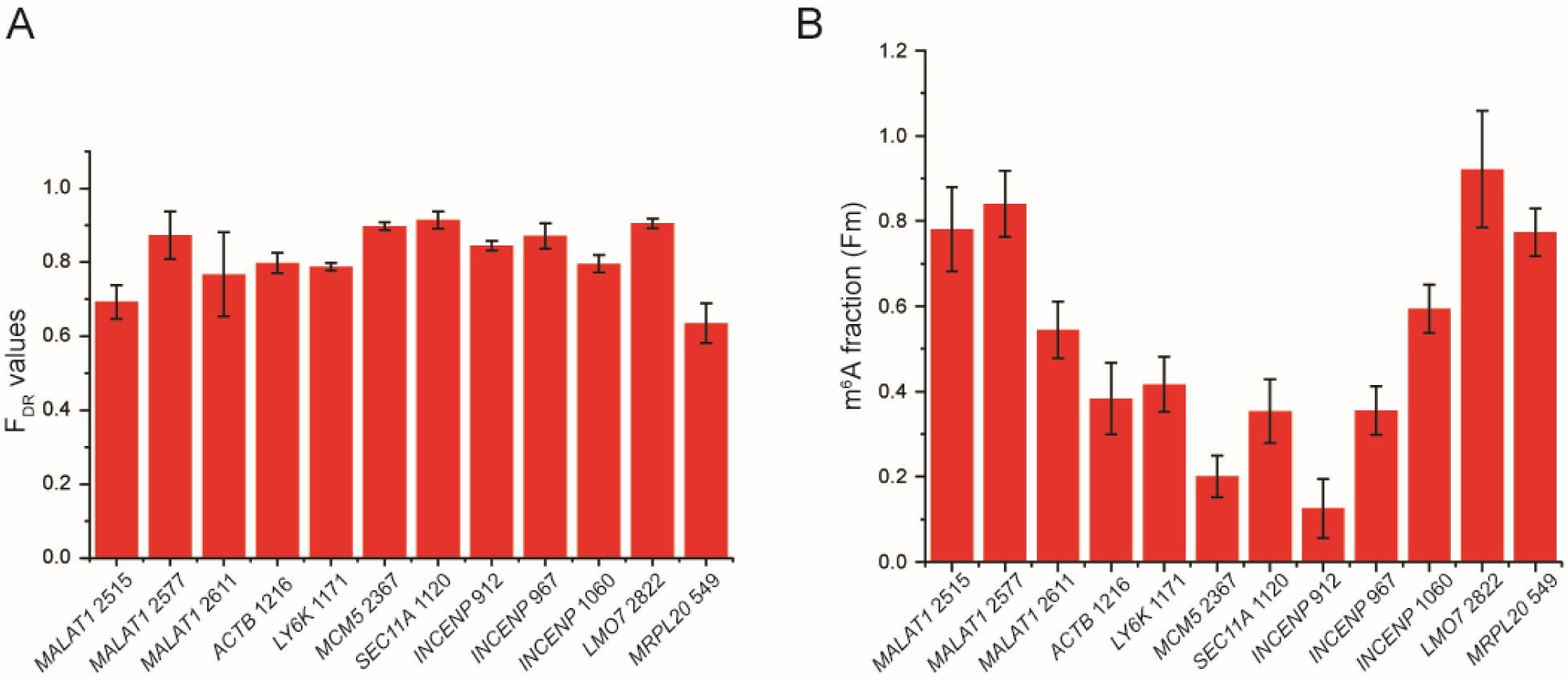
Determination of m^6^A fraction of endogenous sites. (*A*) The cleavage efficiencies (*F*_*DR*_) of the *in vitro* transcribed RNA by 60-mer DR as determined by RT and qPCR. (*B*) Determined m^6^A modification fractions of the 12 endogenous sites. All error bars report mean ± s.d. for at least 3 biological replicates.

The methylation fractions of the endogenous sites were determined to range from 0.20 to 1.0 (Fig. 5B). Notably, four of the targets (*MALAT1* 2515, *MALAT1* 2577, and *MALAT1* 2611 and *ACTB* 1216) were previously measured using the SCARLET assay (Liu et al. 2013), and our results show comparable methylation fractions. While the generally consistent results between our methods and SCARLET assay help validate our assay, we did notice that values measured in our assay are slightly higher than those from SCARLET. One possible explanation for this slight variation can be the splint ligation step used in the SCARLET assay, in which the DNA oligo needs to be ligated to the RNase H cleaved RNA carrying either unmodified A or m^6^A at the 5’ end (Liu et al. 2013). It is possible that the splint ligation is less efficient for the m^6^A containing RNA, and therefore, underestimates the m^6^A fraction in SCARLET assay.

### Potential limitations and other considerations of the method

In summary, we have established a method for quantifying the absolute methylation fraction of potential m^6^A sites of specific transcripts using deoxyribozyme digestion, expanding the toolkit for site-specific quantification of m^6^A. As the VMC10 DR selectively cleaves the unmodified A, it can potentially be used to discriminate other modifications, such as m^1^A (Tserovski et al. 2016). We, therefore, expect the deoxyribozyme-based quantification method can be easily applied to site-specific absolute quantifications of other RNA modifications.

While this method is easy to implement, there are several limitations that need to be considered. Firstly, the assay utilizes VMC10 DR, which has high cleavage efficiencies only on DGACH sequences, limiting its application on a subset of m^6^A sites with the DAACH sequences (Sednev et al. 2018). Secondly, DR digestion efficiency varies among different sequences. Although low DR cleavage efficiency can be corrected by determining *F*_*DR*_ for each modification site of interest using *in vitro* transcribed RNA, low DR efficiency can lead to less accurate quantification due to two reasons. (1) A higher digestion efficiency leads to a larger ΔΔ*Ct* that reduces the measurement variation by qPCR. Conversely, low digestion efficiency will make the ΔΔ*Ct* too small to be accurately detected by qPCR. (2) The <5% cleavage efficiency on the modified RNA can lead to underestimation of the m^6^A fraction, and the percentage of underestimation depends on *F*_*DR*_ (Supplemental Discussion and Supplemental Fig. S1). A lower *F*_*DR*_ will result in a larger underestimation. When *F*_*DR*_ is 50%, a 5% cleavage of the modified RNA will result in a 10% underestimation of the m^6^A. Finally, the presence of a nearby modified nucleotide may affect the DR cleavage efficiency.

To test the effect of the nearby modifications, we designed synthetic RNA containing a nearby m^6^A, m^1^A or ψ and measured their effects on cleavage efficiency of DR by PAGE analysis (Figure 6, Supplemental Figure S6). The cleavage efficiency was unaffected by the presence of m^6^A modification 2 and 4 nt upstream and downstream of the target site, suggesting that the method can be used to quantify m^6^A fraction in RNA that contain m^6^A modifications in clusters. Furthermore, ψ modification had a very minimal decrease in the cleavage efficiency of DR 2 nt away from target site and had no effect on the cleavage when present 4 nt away from the target site. Finally, m^1^A modification, which can affect the Watson-Crick base pairing, significantly decreased the cleavage efficiency at 2 nt away from the target site and moderately decreased the cleavage efficiency at 4 nt away from the target site. Overall, the results indicate that other nearby RNA modifications that do not affect the base pairing with the DR are not likely to affect the DR cleavage efficiency even when placed as close as only 2 nt away from the target site. However, nearby RNA modifications that weaken the base pairing with the DR will have a larger effect on the DR activity, but the effect decreases when the modification is more distal from the target site. To improve the accuracy of the measurement, there are also a few factors to note. Firstly, for the synthetic RNA, we observed equal quality of *F*_*m*_ estimation using samples treated with dDR or samples lacking any DR as a negative control (Fig. 2C; Supplemental Fig. S5). Nevertheless, we still recommend using dDR treated sample as a negative control, because it corrects for potential changes in the RNA secondary structure caused by DR binding that can affect RT efficiency. Secondly, we recommend using 60-mer DR for quantification, as 60-mer DR overall has higher digestion efficiencies of unmethylated RNAs potentially due to a higher hybridization efficiency. Thirdly, the quality of the primers used for RT and qPCR should be verified by performing calibration curves. Finally, we noticed that the largest source of variability in measurements originates from the RT step (comparing error bars in Figure 3B and C). We, thereby, recommend performing multiple RT reactions for each DR treated sample to reduce measurement error.

**FIGURE 6.**
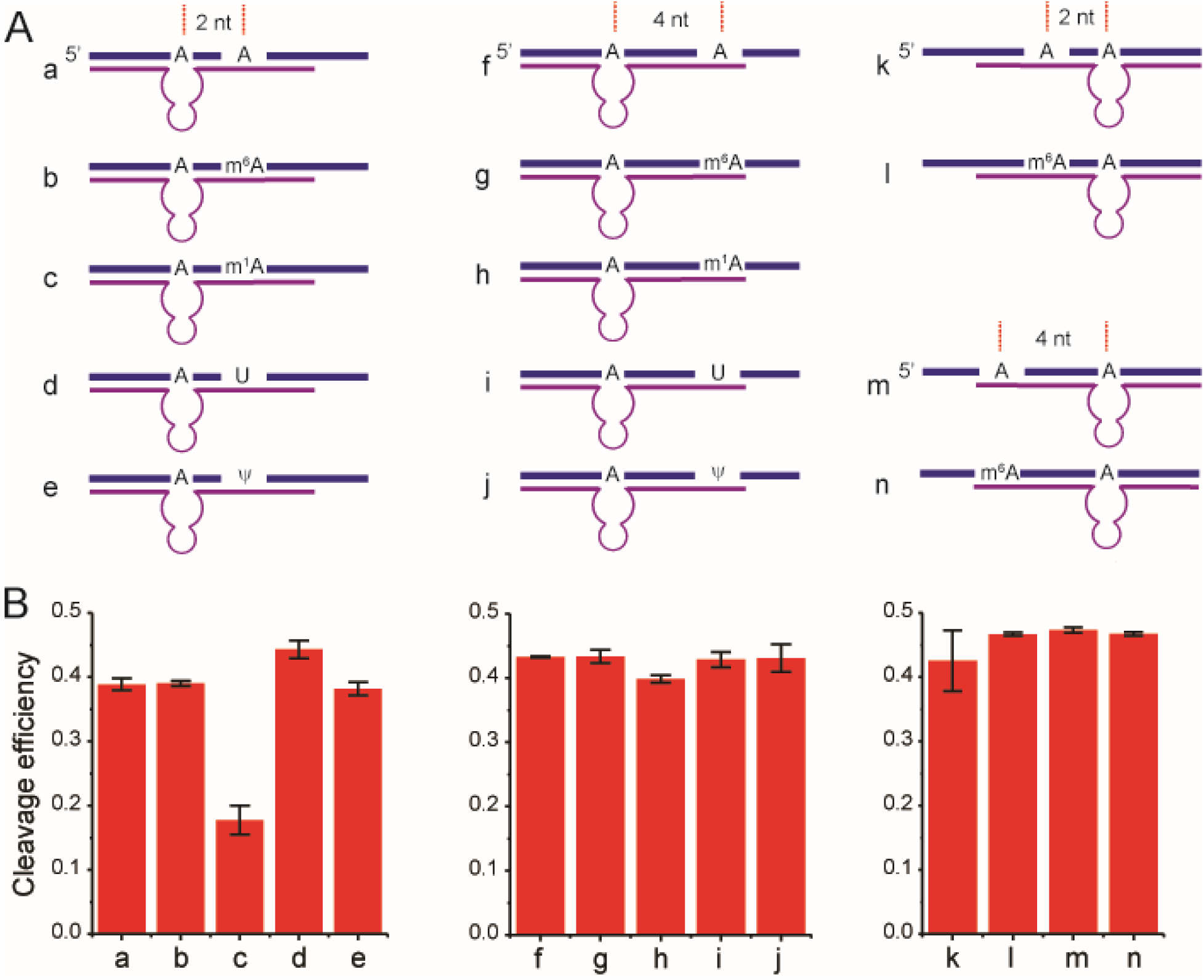
The effects of nearby RNA modifications on cleavage efficiency (*F*_*DR*_) of DR. (*A*) Scheme of 35-nt synthetic RNA containing m^6^A, m^1^A, and ψ modifications. (*B*) Bar plot of the cleavage efficiencies of synthetic RNAs as quantified from PAGE. Error bars indicate mean ± s.d. for 3 independent DR cleavage reactions.

## MATERIALS AND METHODS

### Cell culture and RNA extraction

HeLa cells were cultured in DMEM medium (Gibco) supplemented with 10% (v/v) Fetal Bovine Serum (Thermo Fisher Scientific). The cells were grown at 37 °C under humidified conditions with 5% CO_2_. The total RNA was extracted using RNeasy Mini Kit (Qiagen) according to the manufacturer’s instructions.

### *In vitro* transcription of endogenous RNA target fragments

The dsDNA templates for *in vitro* transcription were prepared by PCR with primers (Integrated DNA Technologies) that contain T7 promoter sequence and cDNA generated from total HeLa RNA. 1 μg of dsDNA templates were added to 100 μl reactions containing final concentrations of 2.5 mM each rNTP (New England Biolabs), 1x T7 Polymerase reaction buffer (New England Biolabs), 4 mM MgCl_2_, 0.5 U/μl SUPERase-In RNase Inhibitor (Thermo Fisher Scientific), and 14 U/μl T7 polymerase (a kind gift from Dr. D. Bishop’s Group). The reactions were incubated at 37 °C for 1 hour. The transcript products were treated with DNase I recombinant (Roche) at 37 °C for 30 min and purified by Phenol-chloroform extraction and ethanol precipitation. Primers are listed in Supplemental Table S2.

### *In vitro* transcription of 0% and 100% methylated GB RNA

A gene block containing 460 nt random sequence with 51% GC content and one adenosine was purchased from Genewiz. The gene block sequence is listed in Supplemental Table S1. The dsDNA template was amplified with primers containing an upstream T7 promoter sequence (Table S2). The *in vitro* reactions were carried out in the same conditions as for endogenous RNA targets, except that N^6^-methyladenosine-5’-triphosphate (Trilink Biotechnologies) was used instead of rATP for the generation of 100% methylated RNA. The transcript products were treated with DNase I recombinant (Roche) at 37 °C for 30 min and purified by phenol-chloroform extraction, 7% denaturing polyacrylamide gel, and ethanol precipitation.

### Synthesis of RNA oligo

Unmodified phosphoramidites were purchased from Glen Research. Phosphoramidite of *N*^6^ - methyladenosine was synthesized by following previously published procedure (Dai et al. 2007). RNA oligos were synthesized using Expedite DNA synthesizer at 1 umol scale. After deprotection, RNA oligos were purified by PAGE.

### Deoxyribozyme digestion

Total RNA (500 ng-2 μg) or *in vitro* transcribed RNA fragments (50 nM) were mixed with 55.6 μM of either DR or dDR (Integrated DNA Technologies), 55.6 mM Tris-HCl (pH 7.5), and 166.7 mM NaCl in a final volume of 9 μl. The annealing of DR to the target site was facilitated by 5 min incubation at 95 °C, followed by slow cooling to room temperature. After annealing, 1 μl of 200 mM MgCl_2_ was added to each reaction and incubated at 37 °C for 12 hours. The final concentrations of the reagents in the incubation buffer are 50 μM of DR, 50 mM Tris-HCl (pH 7.5), and 150 mM NaCl, and 20 mM MgCl_2_ in 10 μl reaction. The DR treatment of 35-40 nt *MALAT1* 2515, *MALAT1* 2577, and *ACTB* 1216 was carried out following the same protocol, except with step-wise cooling (95 °C for 5 min and 25 °C for 10 min) instead of slow cooling. To remove the DR after the digestion, 1.33 μl of 10x TURBO DNase buffer and 2 μl of TURBO DNase (Thermo Fisher Scientific) were added to the 10 μl DR reactions. The samples were incubated at 37 °C for 2 hours. Subsequently, the DNase enzyme was inactivated by addition of EDTA (pH 7.5) to 15 mM final concentration and incubated at 75 °C for 10 min. The DR sequences are listed in the Supplemental Table S3.

### RNA digestion analyzed by PAGE

DR digestion reactions containing 50 ng to 100 ng of *in vitro* transcribed RNA or 35-40 nt synthetic RNAs were run on either 7% or 15% denaturing (7 M urea) polyacrylamide gel electrophoresis (PAGE), respectively. The gels were stained with SYBR Green II RNA Gel Stain (Thermo Fisher Scientific) for 10 min and imaged with ChemiDoc™ Imaging System (Bio-Rad). The cleavage efficiencies were analyzed with ImageJ using intensities of bands corresponding to the full-length RNA and the longer cleaved product.

### Reverse transcription

For each DR treated sample, separate reverse transcription (RT) reactions were performed with gene-specific reverse primers for the m^6^A region and internal reference site. Due to the presence of excess EDTA after DNase inactivation, the reactions were either significantly diluted or extra MgCl_2_ was added for maximum reverse transcriptase activity. The RNA was denatured at 70 °C for 5 min and then added to freshly prepared RT buffer with final concentration of 1 mM dNTPs (Thermo Fisher Scientific), 10% DMSO (Thermo Fisher Scientific), 10 mM DTT (Sigma-Aldrich), 250 nM of gene-specific reverse primer (Integrated DNA Technologies), and 20-fold dilution of reverse transcriptase from iScript cDNA Synthesis Kit (Bio-Rad). The reactions were incubated at 25 °C for 5 min, at 46 °C for 20 min, and heat-inactivated at 95 °C for 1 min. All primers are listed in Supplemental Table S4.

### qPCR

1 μl of cDNA was added into reaction mixture, containing 250 nM of each forward and reverse primers and 1x SsoAdvanced™ Universal SYBR^®^ Green Supermix (Bio-Rad) in a final volume of 20 μl. The qPCR reactions were performed with CFX real-time PCR system (Bio-Rad), using pre-incubation of 95 °C for 30 s, followed by 40 cycles of 95 °C for 10 s and 60 °C for 30 s. The reactions were then subjected to melting curve analysis: 95 °C for 10 s, 65 °C for 5 s increment by 0.5 °C to 95 °C for 5 s. The data was analyzed with the supporting Bio-Rad CFX Maestro software. All primers are listed in Supplemental Table S4. All error bars in the figures are mean ± standard deviation (s.d.) of multiple biological replicates. For *in vitro* prepared GB RNA with different input m6A fraction, biological replicates are defined as independently mixed GB RNA samples. For the cases of endogenous mRNAs, biological replicates are defined as independently extracted total RNA samples. The m^6^A fraction calculated for each biological replicate is from the average values of multiple technical replicates defined by independently performed RT reactions for each RNA sample.

## ACKNOWLEDGMENTS

We thank Dr. C He and Dr. T Pan for useful discussion. Dr. A Driouchi for comments on the manuscript. J Fei acknowledges the support from the Searle Scholars Program, the NIH Director’s New Innovator Award (1DP2GM128185-01), and the pilot grant from the NIH Center for the Dynamic RNA Epitranscriptomes at The University of Chicago.

## AUTHOR CONTRIBUTIONS

M.B. J.Z. and J.F. designed the experiments. M.B., J.Z., and E.H. performed the experiments. M.B. analyzed the data. Q.D. synthesized the 35-40 mer synthetic oligonucleotides. M.B., J.Z. and J.F. wrote the manuscript.

## REFERENCES

Boccaletto P, Machnicka MA, Purta E, Piatkowski P, Baginski B, Wirecki TK, de Crécy-Lagard V, Ross R, Limbach PA, Kotter A, et al. 2018. MODOMICS: a database of RNA modification pathways. 2017 update. Nucleic Acids Res 46: D303–D307.

Chen K, Lu Z, Wang X, Fu Y, Luo G-Z, Liu N, Han D, Dominissini D, Dai Q, Pan T, et al. 2015. High-Resolution *N*^6^ -Methyladenosine (m^6^ A) Map Using Photo-Crosslinking-Assisted m^6^ A Sequencing. Angew Chem Int Ed Engl 127: 1607–1610.

Dai Q, Fong R, Saikia M, Stephenson D, Yu Y, Pan T, Piccirilli JA. 2007. Identification of recognition residues for ligation-based detection and quantitation of pseudouridine and N6-methyladenosine. Nucleic Acids Res 35: 6322–6329.

Dominissini D, Moshitch-Moshkovitz S, Schwartz S, Salmon-Divon M, Ungar L, Osenberg S, Cesarkas K, Jacob-Hirsch J, Amariglio N, Kupiec M, et al. 2012. Topology of the human and mouse m6A RNA methylomes revealed by m6A-seq. Nature 485: 201–206.

Garcia-Campos MA, Edelheit S, Toth U, Safra M, Shachar R, Viukov S, Winkler R, Nir R, Lasman L, Brandis A, et al. 2019. Deciphering the “m6A Code” via Antibody-Independent Quantitative Profiling. Cell.

Harada K, Orgel LE. 1993. Unexpected substrate specificity of T4 DNA ligase revealed by in vitro selection. Nucleic Acids Res 21: 2287–2291.

Harcourt EM, Ehrenschwender T, Batista PJ, Chang HY, Kool ET. 2013. Identification of a selective polymerase enables detection of N(6)-methyladenosine in RNA. J Am Chem Soc 135: 19079–19082.

Helm M, Motorin Y. 2017. Detecting RNA modifications in the epitranscriptome: predict and validate. Nat Rev Genet 18: 275–291.

Jia G, Fu Y, Zhao X, Dai Q, Zheng G, Yang Y, Yi C, Lindahl T, Pan T, Yang Y-G, et al. 2011. N6-methyladenosine in nuclear RNA is a major substrate of the obesity-associated FTO. Nat Chem Biol 7: 885–887.

Kasowitz SD, Ma J, Anderson SJ, Leu NA, Xu Y, Gregory BD, Schultz RM, Wang PJ. 2018. Nuclear m6A reader YTHDC1 regulates alternative polyadenylation and splicing during mouse oocyte development. PLoS Genet 14: e1007412.

Ke S, Pandya-Jones A, Saito Y, Fak JJ, Vågbø CB, Geula S, Hanna JH, Black DL, Darnell JE, Darnell RB. 2017. m6A mRNA modifications are deposited in nascent pre-mRNA and are not required for splicing but do specify cytoplasmic turnover. Genes Dev 31: 990–1006.

Lei Z, Yi C. 2017. A Radiolabeling-Free, qPCR-Based Method for Locus-Specific Pseudouridine Detection. Angew Chem Int Ed Engl 56: 14878–14882.

Linder B, Grozhik AV, Olarerin-George AO, Meydan C, Mason CE, Jaffrey SR. 2015. Single-nucleotide-resolution mapping of m6A and m6Am throughout the transcriptome. Nat Methods 12: 767–772.

Liu J, Yue Y, Han D, Wang X, Fu Y, Zhang L, Jia G, Yu M, Lu Z, Deng X, et al. 2014. A METTL3-METTL14 complex mediates mammalian nuclear RNA N6-adenosine methylation. Nat Chem Biol 10: 93–95.

Liu N, Dai Q, Zheng G, He C, Parisien M, Pan T. 2015. N(6)-methyladenosine-dependent RNA structural switches regulate RNA-protein interactions. Nature 518: 560–564.

Liu N, Parisien M, Dai Q, Zheng G, He C, Pan T. 2013. Probing N6-methyladenosine RNA modification status at single nucleotide resolution in mRNA and long noncoding RNA. RNA 19: 1848–1856.

Liu W, Yan J, Zhang Z, Pian H, Liu C, Li Z. 2018. Identification of a selective DNA ligase for accurate recognition and ultrasensitive quantification of N6-methyladenosine in RNA at one-nucleotide resolution. Chem Sci 9: 3354–3359.

Maity A, Das B. 2016. N6-methyladenosine modification in mRNA: machinery, function and implications for health and diseases. FEBS J 283: 1607–1630.

Meyer KD, Saletore Y, Zumbo P, Elemento O, Mason CE, Jaffrey SR. 2012. Comprehensive analysis of mRNA methylation reveals enrichment in 3’ UTRs and near stop codons. Cell 149: 1635–1646.

Nachtergaele S, He C. 2018. Chemical Modifications in the Life of an mRNA Transcript. Annu Rev Genet 52: 349–372.

Potapov V, Ong JL, Langhorst BW, Bilotti K, Cahoon D, Canton B, Knight TF, Evans TC, Lohman GJS. 2018. A single-molecule sequencing assay for the comprehensive profiling of T4 DNA ligase fidelity and bias during DNA end-joining. Nucleic Acids Res 46: e79.

Santoro SW, Joyce GF. 1997. A general purpose RNA-cleaving DNA enzyme. Proc Natl Acad Sci USA 94: 4262–4266.

Sednev MV, Mykhailiuk V, Choudhury P, Halang J, Sloan KE, Bohnsack MT, Höbartner C. 2018. *n*^6^ -methyladenosine-sensitive rna-cleaving deoxyribozymes. Angew Chem Int Ed Engl 130: 15337–15341.

Shi H, Wei J, He C. 2019. Where, When, and How: Context-Dependent Functions of RNA Methylation Writers, Readers, and Erasers. Mol Cell 74: 640–650.

Tserovski L, Marchand V, Hauenschild R, Blanloeil-Oillo F, Helm M, Motorin Y. 2016. High-throughput sequencing for 1-methyladenosine (m(1)A) mapping in RNA. Methods 107: 110–121.

Wang S, Wang J, Zhang X, Fu B, Song Y, Ma P, Gu K, Zhou X, Zhang X, Tian T, et al. 2016. N^6^ -Methyladenine hinders RNA- and DNA-directed DNA synthesis: application in human rRNA methylation analysis of clinical specimens. Chem Sci 7: 1440–1446.

Warda AS, Kretschmer J, Hackert P, Lenz C, Urlaub H, Höbartner C, Sloan KE, Bohnsack MT. 2017. Human METTL16 is a N6-methyladenosine (m6A) methyltransferase that targets pre-mRNAs and various non-coding RNAs. EMBO Rep 18: 2004–2014.

Xiao Y, Wang Y, Tang Q, Wei L, Zhang X, Jia G. 2018. An Elongation- and Ligation-Based qPCR Amplification Method for the Radiolabeling-Free Detection of Locus-Specific N6-Methyladenosine Modification. Angew Chem Int Ed Engl 57: 15995–16000.

Zhang Z, Chen L-Q, Zhao Y-L, Yang C-G, Roundtree IA, Zhang Z, Ren J, Xie W, He C, Luo G-Z. 2019. Single-base mapping of m^6^ A by an antibody-independent method. Sci Adv 5: eaax0250.

Zheng G, Dahl JA, Niu Y, Fedorcsak P, Huang C-M, Li CJ, Vågbø CB, Shi Y, Wang W-L, Song S-H, et al. 2013. ALKBH5 is a mammalian RNA demethylase that impacts RNA metabolism and mouse fertility. Mol Cell 49: 18–29.

