## Supplemental Information for "Deoxyribozyme-based Method for Site-specific Absolute Quantification of N^6^-methyladenosine Modification Fraction"

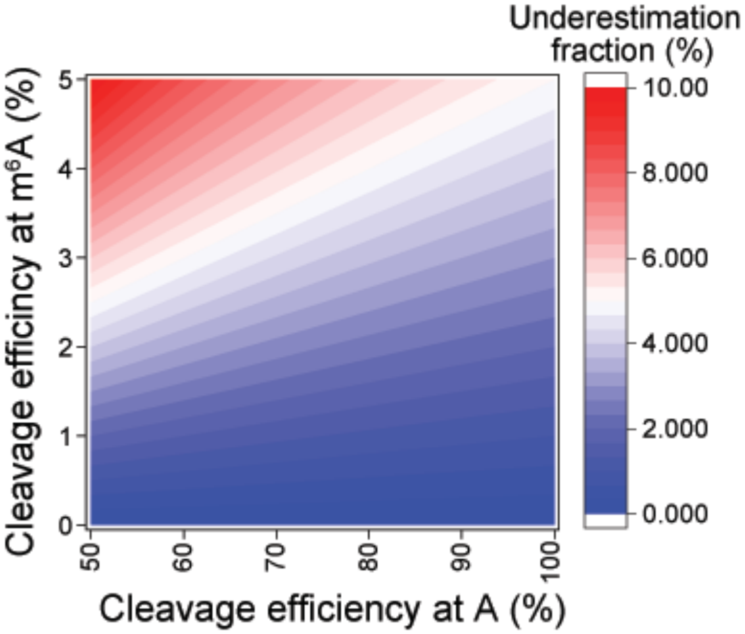


**FIGURE S1**. Percentage of underestimation of m$^6$A fraction due to cleavage of DR on m$^6$A sequence. The trace amount of cleavage of DR at m$^6$A containing sequence will cause false negative signal in the cleavage reaction and underestimation of the m$^6$A percentage. Considering DR cleavage efficiencies of the unmethylated A and m$^6$A sequence are $F_{DR}$ and $F'_{DR}$ respectively, and the true m$^6$A fraction is $F'_m$,

$$(1 - F'_m)F_{DR} + F_m{}'F'_{DR} = 1 - 2^{-\Delta\Delta Ct} \ ,$$

where $\Delta\Delta Ct$ is determined in Eq (2). Comparing $F'_m$ to $F_m$ from Eq (7) determined without the correction of $F'_{DR}$, the percentage of underestimation is

$$\frac{F'_m - F_m}{F'_m} = \frac{F'_{DR}}{F_{DR}} \ .$$

The heat map shows the percentage of underestimation as a function of $F_{DR}$ and $F'_{DR}$, which increases when $F_{DR}$ decreases, assuming $F'_{DR}$ remains lower than 5%. When $F_{DR}$ is 50% and $F'_{DR}$ is 5%, the error in m6A fraction is 10%.

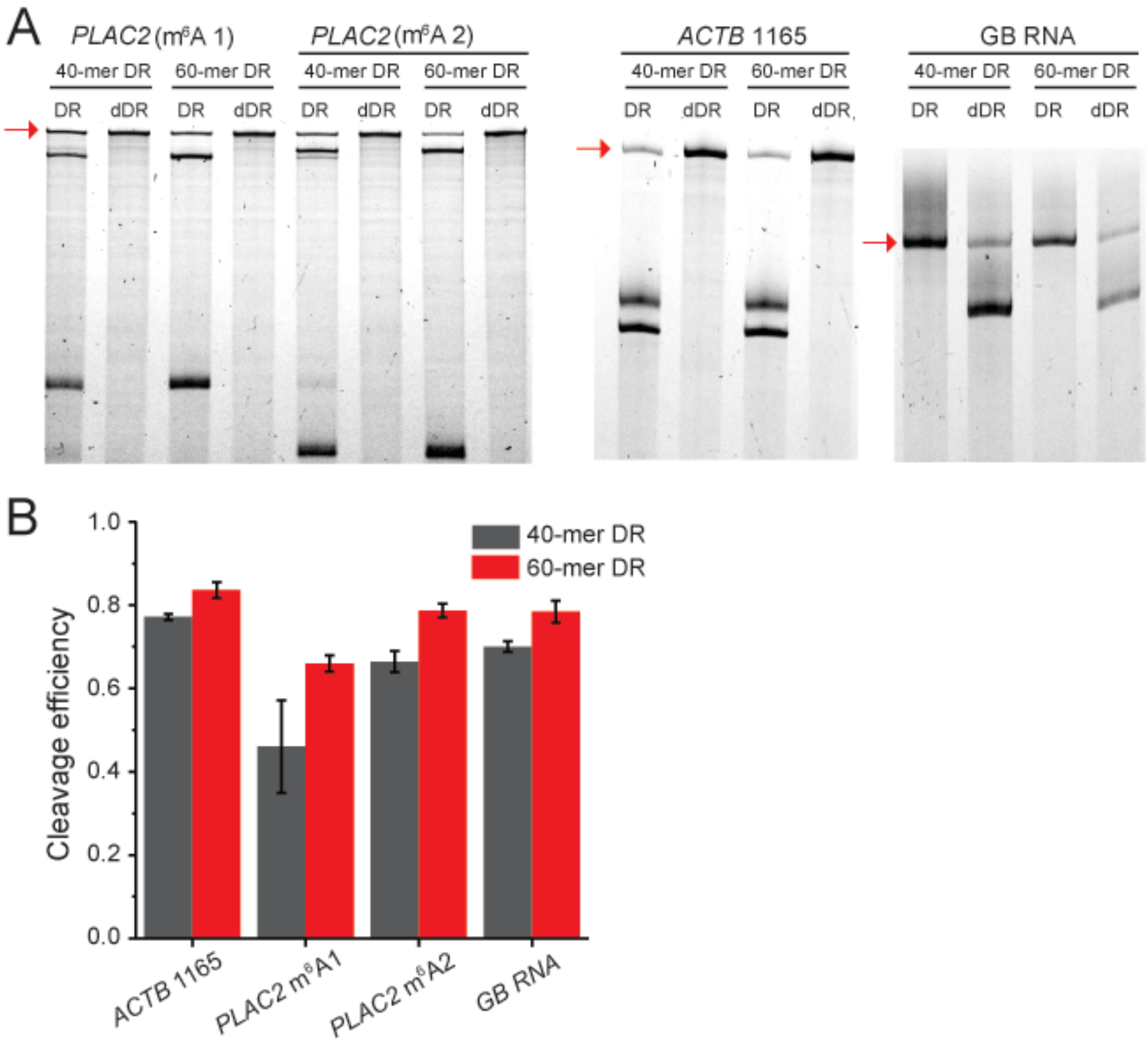


**FIGURE S2**. Cleavage efficiencies ($F_{DR}$) of unmodified *in vitro* transcribed RNA by 40-mer and 60-mer DR and dDR. (*A*) PAGE showing the DR cleavage of *PLAC2* m$^6$A1 and m$^6$A2 sites, *ACTB* 1165, and GB RNA by 40-mer and 60-mer DR. (*B*) Bar plot of the cleavage efficiencies of *PLAC2* m$^6$A1 and m$^6$A2, *ACTB* 1165, and the GB RNA by 40-mer and 60-mer DR as quantified from (*A*)-(*C*). Error bars indicate mean ± s.d. for 3 independent DR cleavage reactions. Red arrows point to full-length uncleaved RNA fragments.

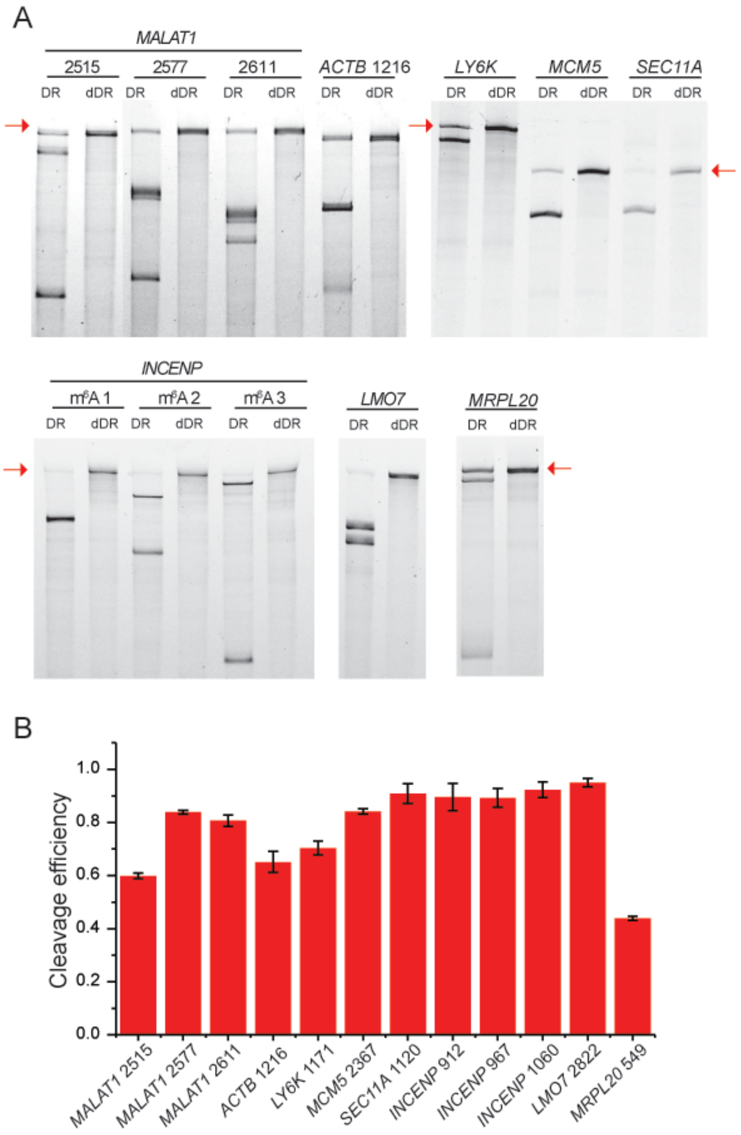


**FIGURE S3**. Cleavage efficiencies ($F_{DR}$) of unmodified *in vitro* transcribed RNA by 60-mer DR and dDR. (*A*) PAGE showing the DR cleavage of the seven endogenous targets: *MALAT1* 2515, 2577, and 2611, *ACTB* 1216, *LY6K* 1171, *MCM5* 2367, *SEC11A* 1120, *INCENP* 912, 967 and 1060, *LMO7* 2822, and *MRPL20* 549. (*B*) Bar plot of the cleavage efficiencies of endogenous targets as quantified from (*A*). Error bars indicate mean ± s.d. for 3 DR cleavage reactions. Red arrows point to full-length uncleaved RNA fragments.

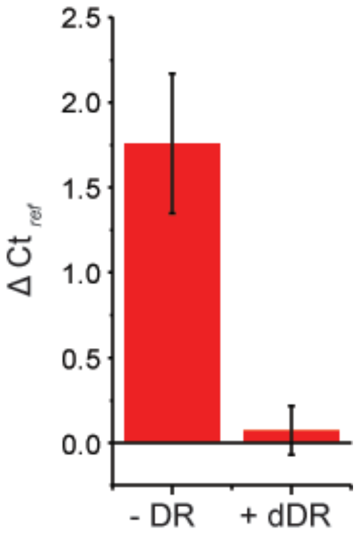


**FIGURE S4**. Negative control without DR and with non-functional version of DR (dDR). When the negative control does not contain DR, there is a consistent difference between $Ct_{+DR-ref}$ and $Ct_{-DR-ref}$ ($\Delta\ Ct_{ref}$), with $Ct_{-DR-ref}$ being larger. Use of dDR in the negative control eliminates $\Delta\ Ct_{ref}$. Data comes from quantification of GB RNA with 0.0, 0.2, 0.4, 0.6, 0.8, and 1.0 $m^6A$ fraction input. Error bars indicate mean ± s.d.

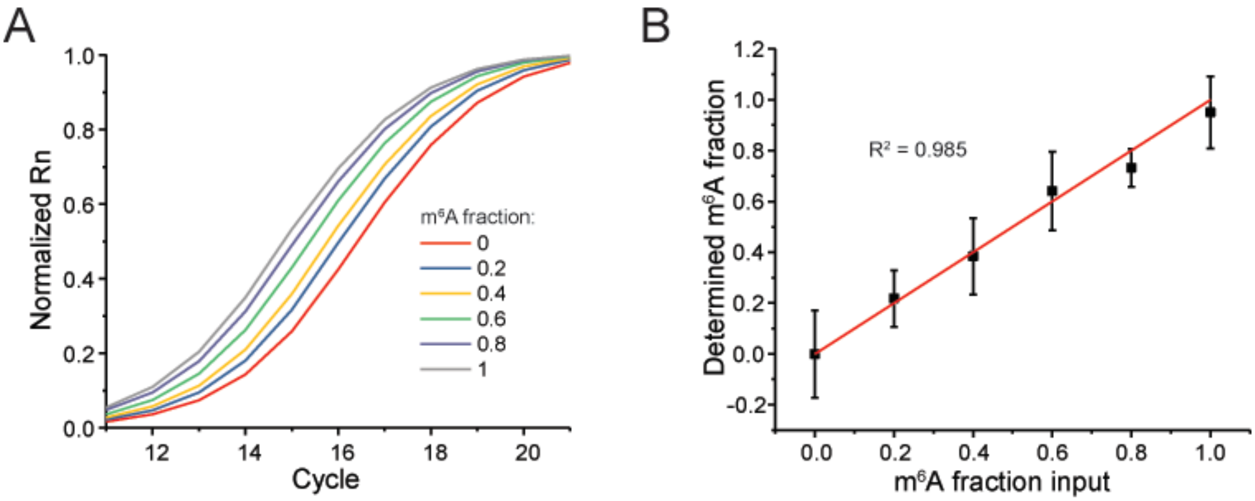


**FIGURE S5**. Validation of the method for absolute quantification of $m^6A$ fraction of the GB RNA without dDR. (*A*) Normalized real-time fluorescence amplification curves for the DR cleaved synthetic RNAs with primers amplifying the $m^6A$ site. (*B*) Estimated modification fraction as a function of input $m^6A$ fraction for the GB RNA. Error bars indicate mean ± s.d. for 3 biological replicates.

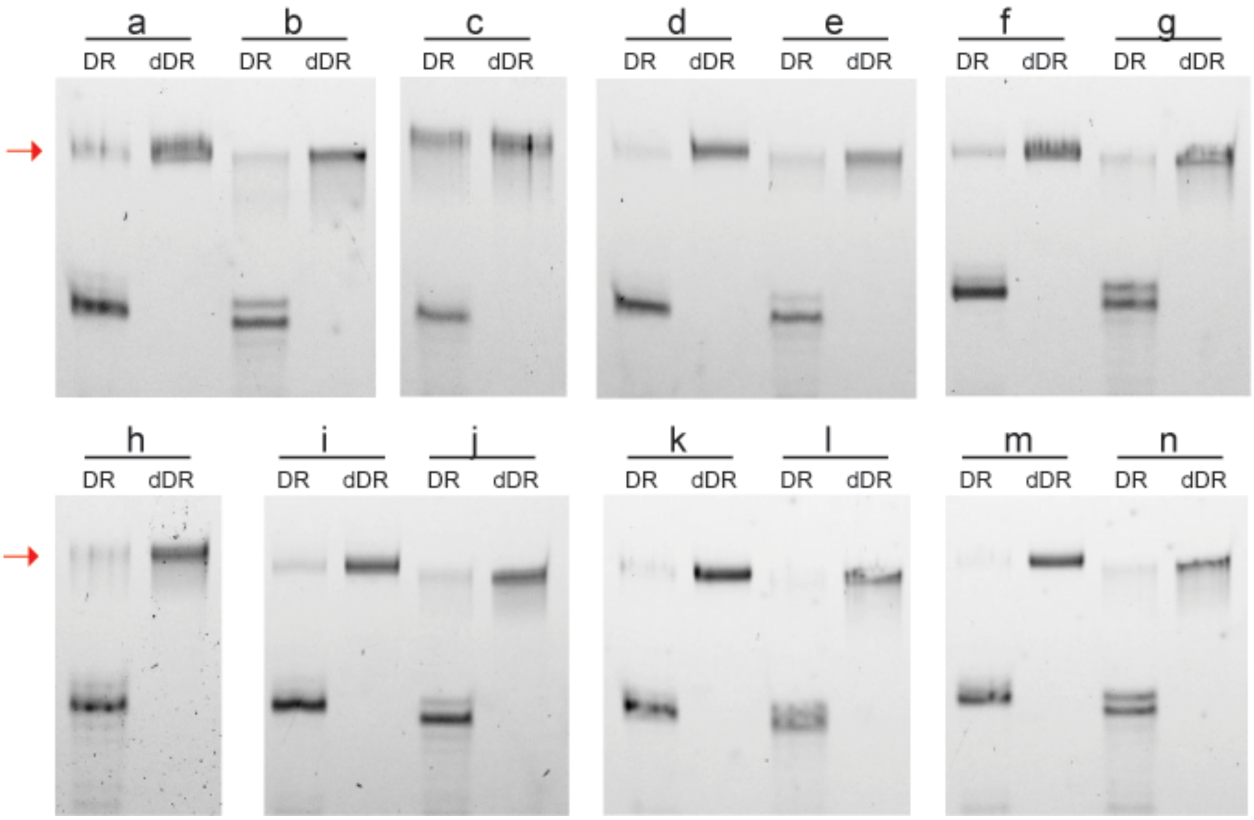


**FIGURE S6**. PAGE showing the DR cleavage efficiency in absence and presence of nearby $m^6A$, $m^1A$, and ψ modifications. Labels a-n correspond to 35-nt synthetic RNAs shown in Figure 6A and listed in Supplemental Table S1. Red arrows point to full-length uncleaved RNA.

**Table S1**: Synthetic DNA and RNA sequences.

| Description | Sequence |
|---|---|
| GB DNA template | GGTTGCGTTGGGTGTTCCTGTTTCTTTTGGCCTTTGTCTCTGTTTCT TTCCTTTCTCCTCCTTGTCGTGTCTGTTCGTTTCGTCGCTTTCCTCC TTCCTTGTTCTCGCTCGGACTCTTCTGGGCTCTTTCTGCGTTCGCCC TTCTTGTTCTCCCTTCTCTGTGGGTCCTGTTCTTGTGCTGGTTGTGC TCCCTCCTCTTGGTGCTCCTCCTTTCTGTGGCTGCGCTGGTGTTTCT TTCTCTCGGCTGCTCTGTTTGTTGGGTCTTTGTTGTGTGTTGTTGTT CTTGTGTGCTGCGTTTTGGTGGTGTCGGTTCTGTGCTGTCTTTCGGC CTGTCGTTTTCCTTCTCGTGTTCCGTCCTGTTTTGCGTGTCTCTCCCT GTGTTCCCGCTTTCCGTGTTGGCTGTGCTTGGTGTCTTTCGCTTGTTG GTTGGTCTCCTGTCTCCTGTGCTCGTCGGTCTTGTGG |
| 41-nt synthetic *ACTB* 1216 | CGCAAAUGCUUCUAGGCGGACUAUGACUUAGUUGCGUUACU |
| 41-nt synthetic *MALAT1* 2515 | AGUUUGAAAAAUGUGAAGGACUUUCGUAACGGAAGUAAUUU |
| 32-nt synthetic *MALAT1* 2577 | AACUUAAUGUUUUUGCAUUGGACUUUGAGUUA |
| (a) 35-nt synthetic A 2-nt A control 1 | GCCUUGUUCUCGCUCGGACUAUUCUGGGCUCUUUC |
| (b) 35-nt synthetic A 2-nt $m^6$A | GCCUUGUUCUCGCUCGGACU$m^6$AUUCUGGGCUCUUUC |
| (c) 35-nt synthetic A 2-nt $m^1$A | GCCUUGUUCUCGCUCGGACU$m^1$AUUCUGGGCUCUUUC |
| (d) 35-nt synthetic A 2-nt U control | GCCUUGUUCUCGCUCGGACUUUUCUGGGCUCUUUC |
| (e) 35-nt synthetic A 2-nt ψ | GCCUUGUUCUCGCUCGGACUψUUCUGGGCUCUUUC |
| (f) 35-nt synthetic A 4-nt A control 1 | GCCUUGUUCUCGCUCGGACUCUACUGGGCUCUUUC |
| (g) 35-nt synthetic A4-nt $m^6$A | GCCUUGUUCUCGCUCGGACUCU$m^6$ACUGGGCUCUUUC |
| (h) 35-nt synthetic A 4-nt $m^1$A | GCCUUGUUCUCGCUCGGACUCU$m^1$ACUGGGCUCUUUC |
| (i) 35-nt synthetic A 4-nt U control | GCCUUGUUCUCGCUCGGACUCUUCUGGGCUCUUUC |

| | |
|---|---|
| (j) 35-nt synthetic A 4-nt ψ | GCCUUGUUCUCGCUCGGACUCUψCUGGGCUCUUUC |
| (k) 35-nt synthetic A 2-nt A control 2 | GCCUUGUUCUCGCUAGGACUCUUCUGGGCUCUUUC |
| (l) 35-nt synthetic $m^6A$ 2-nt A | GCCUUGUUCUCGCU$m^6A$GGACUCUUCUGGGCUCUUUC |
| (m) 35-nt synthetic A 4-nt A control 2 | GCCUUGUUCUCGAUCGGACUCUUCUGGGCUCUUUC |
| (n) 35-nt synthetic $m^6A$ 4-nt A | GCCUUGUUCUCG$m^6A$UCGGACUCUUCUGGGCUCUUUC |

**Table S2:** Primers used for generating templates for *in vitro* transcription.

| Description | Sequence |
|---|---|
| Forward primer GB RNA DNA template | TAATACGACTCACTATAGGGTTGCGTTGGGTGTTCCTG |
| Reverse primer for GB RNA DNA template | CCACAAGACCGACGAGCACA |
| Forward primer for *ACTB* DNA template | TAATACGACTCACTATAGGCCAACACAGTGCTGTCTGGC |
| Reverse primer for *ACTB* DNA template | CTGCTGTCACCTTCACCGTTCC |
| Forward primer for *PLAC2* DNA template | TAATACGACTCACTATAGCAAGCAAAGTGAACACGTCG |
| Reverse primer for *PLAC2* DNA template | GTACTGACGTCGGCATCGAT |
| Forward primer for *MALAT1* DNA template | TAATACGACTCACTATAGGCTACTAAAAGGACTGGTGT |
| Reverse primer for *MALAT1* DNA template | TTCACCACCAAATCGTTAGC |
| Forward primer for *LY6K* DNA template | TAATACGACTCACTATAGGCAGGCCATACCACGCAGAAG |
| Reverse primer for *LY6K* DNA template | CCAAGACCCTGGGAAGTCAAA |
| Forward primer for *MCM5* DNA template | TAATACGACTCACTATAGGGAGATGCTGAGCCGCATC |
| Reverse primer for *MCM5* DNA template | CAGCAGGACACTACAGCTCC |
| Forward primer for *SEC11A* DNA template | TAATACGACTCACTATAGGGTCTGTGATTGGTGGAATGG |
| Reverse primer for *SEC11A* DNA template | AAGACTTACGACCACCTCAG |
| Forward primer for *INCENP* DNA template | TAATACGACTCACTATAGATAACCACACCCAGTGCCAG |
| Reverse primer for *INCENP* DNA template | TGCGGACAACACTTTCCTGT |
| Forward primer for *LMO7* DNA template | TAATACGACTCACTATAGGAAATGCTGCAGGACAGGGA |
| Reverse primer for *LMO7* DNA template | TGAGAGCCAAAGGGTCTTGG |
| Forward primer for *MRPL20* DNA template | TAATACGACTCACTATAGGCCGCTACTTTCGGATCCAGG |
| Reverse primer for *MRPL20* DNA template | GGCCATCCCTCATGTCTGTT |

**Table S3**: Deoxyribozyme sequences.

| Description | Sequence |
|---|---|
| GB RNA DR 40-mer | CCCAGAAGAGGGGTCTCCAGCTGGACGTTCGAGCGAGAAC |
| GB RNA dDR 40-mer | CCCAGAAGAGGGGTCTCCTCGTGGATTTCCGAGCGAGAAC |
| GB RNA DR 60-mer | GCAGAAAGAGCCCAGAAGAGGGGTCTCCAGCTGGACGTT CGAGCGAGAACAAGGAAGGAG |
| GB RNA dDR 60-mer | GCAGAAAGAGCCCAGAAGAGGGGTCTCCTCGTGGATTTCC GAGCGAGAACAAGGAAGGAG |
| *PLAC2* m$^6$A 1 DR 40-mer | CCTCTGAGTGGGGTCTCCAGCTGGACGTTACTCCTGCCCC |
| *PLAC2* m$^6$A 2 DR 40-mer | TGGGAAAATGGGGTCTCCAGCTGGACGTTCTGGGCAAGAG |
| *PLAC2* m$^6$A 1 dDR 40-mer | CCTCTGAGTGGGGTCTCCTCGTGGATTTCACTCCTGCCCC |
| *PLAC2* m$^6$A 2 dDR 40-mer | TGGGAAAATGGGGTCTCCTCGTGGATTTCCTGGGCAAGAG |
| *PLAC2* m$^6$A 1 DR 60-mer | AGCGGAAGTGCCTCTGAGTGGGGTCTCCAGCTGGACGTTA CTCCTGCCCCTTCTGTGCTT |
| *PLAC2* m$^6$A 2 DR 60-mer | AAGGTGTGGCTGGGAAAATGGGGTCTCCAGCTGGACGTTC TGGGCAAGAGCGGAAGTGCC |
| *PLAC2* m$^6$A 1 dDR 60-mer | AGCGGAAGTGCCTCTGAGTGGGGTCTCCTCGTGGATTTCA CTCCTGCCCCTTCTGTGCTT |
| *PLAC2* m$^6$A 2 dDR 60-mer | AAGGTGTGGCTGGGAAAATGGGGTCTCCTCGTGGATTTCC TGGGCAAGAGCGGAAGTGCC |
| *ACTB* 1165 DR 40-mer | CTCGTCATACGGGTCTCCAGCTGGACGTTCTGCTTGCTGA |
| *ACTB* 1165 dDR 40-mer | CTCGTCATACGGGTCTCCTCGTGGATTTCCTGCTTGCTGA |
| *ACTB* 1165 DR 60-mer | AGGGGCCGGACTCGTCATACGGGTCTCCAGCTGGACGTTC TGCTTGCTGATCCACATCTG |

| | |
|---|---|
| *ACTB* 1165 dDR 60-mer | AGGGGCCGGACTCGTCATACGGGTCTCCTCGTGGATTTCCT GCTTGCTGATCCACATCTG |
| *ACTB* 1216 DR 60-mer | GTAACGCAACTAAGTCATAGGGGTCTCCAGCTGGACGTTC GCCTAGAAGCATTTGCGGTG |
| *ACTB* 1216 dDR 60-mer | GTAACGCAACTAAGTCATAGGGGTCTCCTCGTGGATTTCC GCCTAGAAGCATTTGCGGTG |
| *MALAT1* 2515 DR 60-mer | AATTACTTCCGTTACGAAAGGGGTCTCCAGCTGGACGTTCT TCACATTTTTCAAACTAAG |
| *MALAT1* 2515 dDR 60-mer | AATTACTTCCGTTACGAAAGGGGTCTCCTCGTGGATTTCCT TCACATTTTTCAAACTAAG |
| *MALAT1* 2577 DR 60-mer | AAAATAATCTTAACTCAAAGGGGTCTCCAGCTGGACGTTC AATGCAAAAACATTAAGTTG |
| *MALAT1* 2577 dDR 60-mer | AAAATAATCTTAACTCAAAGGGGTCTCCTCGTGGATTTCC AATGCAAAAACATTAAGTTG |
| *MALAT1* 2611 DR 60-mer | CAGCTGTCAATTAATGCTAGGGGTCTCCAGCTGGACGTTCT CAGGATTTAAAAAATAATC |
| *MALAT1* 2611 dDR 60-mer | CAGCTGTCAATTAATGCTAGGGGTCTCCTCGTGGATTTCCT CAGGATTTAAAAAATAATC |
| *LY6K* DR 60-mer | GAAGGCTCAGTCTGTGGCAGGGGTCTCCAGCTGGACGTTC CGTGGCTCAAGACAGGCTGA |
| *LY6K* dDR 60-mer | GAAGGCTCAGTCTGTGGCAGGGGTCTCCTCGTGGATTTCC CGTGGCTCAAGACAGGCTGA |
| *MCM5* DR 60-mer | CAGAGGTCCCAGCAACATTGGGGTCTCCAGCTGGACGTTA ATGGCAGGCAGCGGCAGGAG |

| | |
|---|---|
| *MCM5* dDR 60-mer | CAGAGGTCCCAGCAACATTGGGGTCTCCTCGTGGATTTCA ATGGCAGGCAGCGGCAGGAG |
| *SEC11A* DR 60-mer | GCTGCATTTTCATTTACAAGGGGTCTCCAGCTGGACGTTTC TGTAGGCACTTTAGAAGTG |
| *SEC11A* dDR 60-mer | GCTGCATTTTCATTTACAAGGGGTCTCCTCGTGGATTTCTC TGTAGGCACTTTAGAAGTG |
| *INCENP* 912 DR 60-mer | CTTAGACGCAGACCGCCCCGGGGTCTCCAGCTGGACGTTC CGACCCCTTGACCCTTGGGG |
| *INCENP* 912 dDR 60-mer | CTTAGACGCAGACCGCCCCGGGGTCTCCTCGTGGATTTCCC GACCCCTTGACCCTTGGGG |
| *INCENP* 967 DR 60-mer | AATCTGGAAAGGCTGGCGAGGGGTCTCCAGCTGGACGTTC CGTGGGCCAGGGGAGACCTG |
| *INCENP* 967 dDR 60-mer | AATCTGGAAAGGCTGGCGAGGGGTCTCCTCGTGGATTTCC CGTGGGCCAGGGGAGACCTG |
| *INCENP* 1060 DR 60-mer | TGTGCCGCACCGATTGAGAGGGGTCTCCAGCTGGACGTTC GTGCGAGAGCCCGTGGGCGT |
| *INCENP* 1060 dDR 60-mer | TGTGCCGCACCGATTGAGAGGGGTCTCCTCGTGGATTTCC GTGCGAGAGCCCGTGGGCGT |
| *LMO7* DR 60-mer | GAATTTCAGTTGTTACACGGGGGTCTCCAGCTGGACGTTCT CTCTTTTTGCAAAAGTGGT |
| *LMO7* dDR 60-mer | GAATTTCAGTTGTTACACGGGGGTCTCCTCGTGGATTTCCT CTCTTTTTGCAAAAGTGGT |
| *MRPL20* DR 60-mer | CCTAATCAATACAGCAACAGGGGTCTCCAGCTGGACGTTC TCAGTGGTACTGCACCACTC |

| | |
|---|---|
| *MRPL20* dDR 60-mer | CCTAATCAATACAGCAACAGGGGTCTCCTCGTGGATTTCCT CAGTGGTACTGCACCACTC |

**Table S4:** Primers for reverse transcription and qPCR.

| Description | Sequence |
|---|---|
| Forward primer GB RNA m$^6$A region | GGTTGCGTTGGGTGTTCCTG |
| Reverse primer for GB RNA m$^6$A region | GGGAGAACAAGAAGGGCGAA |
| Forward primer for GB RNA control region | CGTCCTGTTTTGCGTGTCTC |
| Reverse primer for GB RNA control region | CCACAAGACCGACGAGCACA |
| Forward primer for *ACTB* m$^6$A region | CCTTCCAGCAGATGTGGATC |
| Reverse primer for *ACTB* m$^6$A region | GCCATGCCAATCTCATCTTG |
| Forward primer for *ACTB* control region | CAGGATGCAGAAGGAGATCAC |
| Reverse primer for *ACTB* control region | CGATCCACACGGAGTACTTG |
| Forward primer for *PLAC2* m$^6$A region | AAGAGAAGCACAGAAGGGGC |
| Reverse primer for *PLAC2* m$^6$A region | ACGGCTTGGGCAAAGGTGTG |
| Forward primer for *PLAC2* control region | CAAGCAAAGTGAACACGTCG |
| Reverse primer for *PLAC2* control region | TCACTTTAACTTGCACTTTACTGC |
| Forward primer for *MALAT1* m$^6$A region | GGCAGAAGGCTTTTGGAAGAGT |
| Reverse primer for *MALAT1* m$^6$A region | CTGGGTCAGCTGTCAATTAATGC |
| Forward primer for *MALAT1* control region | CAGCAGCAGACAGGATTCCA |
| Reverse primer for *MALAT1* control region | TCCTATCTTCACCACGAACTGC |
| Forward primer for *LY6K* m$^6$A region | GGCCTCAGCCTGTCTTGA |
| Reverse primer for *LY6K* m$^6$A region | AATGCAACAGGTGACAACGG |
| Forward primer for *LY6K* control region | TGACTGTGCACCTTTGAGCA |
| Reverse primer for *LY6K* control region | ACCGAGAGAAGGCAATCACG |
| Forward primer for *MCM5* m$^6$A region | TCACTGGACTCATGGACTCG |
| Reverse primer for *MCM5* m$^6$A region | AAGTTCGAGGGCTGCAGT |
| Forward primer for *MCM5* control region | GAGCACAGCATCATCAAGGA |

| | |
|---|---|
| Reverse primer for *MCM5* control region | TGCATGCGATGCTGGATCT |
| Forward primer for *SEC11A* m$^6$A region | CAAAGCCCCCAGTGTTTGTA |
| Reverse primer for *SEC11A* m$^6$A region | CGTGCAGAGCTGCATTTTCAT |
| Forward primer for *SEC11A* control region | CACTCGAGGGGACTTTCAGT |
| Reverse primer for *SEC11A* control region | GGCTTTGGCTCAACCTTTTAAT |
| Forward primer for *INCENP* 912 and 967 m$^6$A region | TGAGCTCCCTGATGGCTACA |
| Reverse primer for *INCENP* 912 and 967 m$^6$A region | CTCCCGCCATGGAGAATCTG |
| Forward primer for *INCENP* 1060 m$^6$A region | CTCCCATCCTGCCGGATAAC |
| Reverse primer for *INCENP* 1060 m$^6$A region | TGGGCTAAGACTTGGGGACT |
| Forward primer for *INCENP* control region | CATCAGTGAGCGCCAGAATG |
| Reverse primer for *INCENP* control region | TGATGTCGGGATGCCCTG |
| Forward primer for *LMO7* m$^6$A region | GAGAGAGTAGAAGAGAAGGG |
| Reverse primer for *LMO7* m$^6$A region | CAAAGAGGCTGGGCTTTGTTC |
| Forward primer for *LMO7* control region | TCACGGAGCACACAAATGGA |
| Reverse primer for *LMO7* control region | TGAGAGCCAAAGGGTCTTGG |
| Forward primer for *MRPL20* m$^6$A region | GGGAAGGAACCTGAAGGCAT |
| Reverse primer for *MRPL20* m$^6$A region | TGCAAATTACTCTGTCTCTTTTCC |
| Forward primer for *MRPL20* control region | CCAAAGCCCGATACCTGAAGA |
| Reverse primer for *MRPL20* control region | GCTCCACCTGGCACTTAACTA |
